# Natural variation in root exudate composition in the genetically structured *Arabidopsis thaliana* in the Iberian Peninsula

**DOI:** 10.1101/2024.09.11.612578

**Authors:** Harihar Jaishree Subrahmaniam, F. Xavier Picó, Thomas Bataillon, Camilla Lind Salomonsen, Marianne Glasius, Bodil K. Ehlers

**Affiliations:** Department of Ecoscience, Aarhus University, 8000 Aarhus C, Denmark; Departamento de Ecología y Evolución, Estación Biológica de Doñana, Consejo Superior de Investigaciones Científicas, 41092 Sevilla, Spain; Department of Molecular Biology and Genetics, Bioinformatics Research Centre, Aarhus University, 8000 Aarhus C, Denmark; Department of Chemistry, Aarhus University, 8000 Aarhus C, Denmark; Universität Hamburg, Institut für Pflanzenwissenschaften und Mikrobiologie, 22609, Hamburg, Germany

**Keywords:** *Arabidopsis thaliana*, drivers of natural variation, genetic clusters, genome-wide association analysis, root exudates, terpenoids

## Abstract

- Plant root exudates are involved in nutrient acquisition, microbial partnerships, and inter-organism signaling. Yet, little is known about the genetic and environmental drivers of root exudate variation at large geographical scales, which may help understand evolutionary trajectories of plants in heterogeneous environments.
- We quantified natural variation in chemical composition of *Arabidopsis thaliana* root exudates in 105 Iberian accessions. We identified up to 373 putative compounds using ultra-high performance liquid chromatography coupled with mass spectrometry. We estimated broad-sense heritability of compounds and conducted a genome-wide association (GWA) study. We associated variation in root exudates to variation in geographic, environmental, life history, and genetic attributes of Iberian accessions.
- Only 25 of 373 compounds exhibited broad-sense heritability values significantly different from zero. GWA analysis identified polymorphisms associated to 12 root exudate compounds and 26 known genes involved in metabolism, defense, signaling, and nutrient transport. The genetic structure influenced root exudate composition involving terpenoids. We detected five terpenoids related to plant defense significantly varying in mean abundances in two genetic clusters.
- Our study provides first insights into the extent of root exudate natural variation at a regional scale depicting a diversified evolutionary trajectory among *A. thaliana* genetic clusters chiefly mediated by terpenoid composition.

## Introduction

Plant root exudates encompass a vast array of primary (e.g. carbohydrates, amino acids, organic acids) and secondary metabolites (e.g. flavonoids, terpenoids, alkaloids) that shape the physical, chemical, and biological properties of the soil (Oburger & Jones, 2018). They also facilitate nutrient cycling and mediate biotic interactions in the rhizosphere, thereby fostering a healthy soil ecosystem (Badri & Vivanco, 2009; Rasmann & Hiltpold, 2022). Despite the ecological relevance of root exudates, several factors, such as stress and developmental status, influence their chemical composition and challenge their quantification. For instance, stress by elevated phosphorus increases pthalic acid in *Cyperus alternifolius* (Duan *et al*., 2020), hydric stress induces various organic acids in *Zea mays* (Song *et al*., 2012), and pathogen infection in *Arabidopsis thaliana* stimulates long-chain fatty acids and amino acids that recruit protective *Pseudomonas* species (Wen *et al*., 2021). Furthermore, development also affects exudate profiles in *A. thaliana* with sugar alcohols decreasing and amino acids increasing over time in early developmental stages (Chaparro *et al*., 2013), whereas young fir trees exudate more carbohydrates and quercetin than older trees, which secrete more lipids and salicylic acids, shifting from nutrient acquisition to defense over development (Chen *et al*., 2023). Given the influence of root exudates on plant-environment interactions and adaptive strategies (Novoplansky, 2019; Williams & de Vries, 2020; Subrahmaniam *et al*., 2023), unravelling the chemistry of root exudates may help decipher the complexity of plant metabolism, but also the ecology of plant communities (Mommer *et al*., 2016; van Dam & Bouwmeester, 2017; McLaughlin *et al*., 2023).

However, our knowledge of natural variation in root exudates composition is rather scarce (Vives-Peris *et al*., 2020; Escolà Casas & Matamoros, 2021; Wang *et al*., 2021). This is a problem because understanding natural variation in plant traits has a paramount importance in different disciplines, as natural variation reflects long-term evolutionary dynamics, can reveal environmental factors driving this variation, and facilitates the exploration of the genetic basis of trait differences (Mitchell-Olds & Schmitt, 2006; Alonso-Blanco *et al*., 2009). One reason for the scarcity of studies on natural variation in plant root exudates has to do with the technical challenges for capturing and analyzing the complex chemical data from root exudates (van Dam & Bouwmeester, 2017; Oburger & Jones, 2018). In fact, few studies have described natural variation in chemical composition of root exudates in various plant species (Micallef *et al*., 2009; Biedrzycki *et al*., 2010; Badri *et al*., 2012; Houshyani *et al*., 2012; Chaparro *et al*., 2013; Fang *et al*., 2013; Strehmel *et al*., 2014; Mönchgesang *et al*., 2016; Kawasaki *et al*., 2018; Liu *et al*., 2020), all of them using low sample sizes (less than 20 accessions in all cases). Interestingly, recent developments in mass spectrometry and nuclear magnetic resonance spectroscopy now enable the characterization and quantification of specific chemical compounds present in root exudates from a large number of samples (Pantigoso *et al*., 2021; Wang *et al*., 2022).

In this study, we took advantage of such recent technical advances, combined with the availability of dense collections of natural accessions exceptionally well-characterized at the ecological, phenotypic, and genomic levels of the annual plant *Arabidopsis thaliana*, to conduct the first regional-scale assessment of natural variation in root exudate composition in plants with a large sample size. We analyzed root exudates from 105 distinct natural accessions of *A. thaliana* from the Iberian Peninsula. The Iberian collection of *A. thaliana* is geographically structured into four differentiated genetic clusters (Picó *et al*., 2008; Castilla *et al*., 2020), reflecting the complexity of the demographic and evolutionary history of this species across the region. Besides, the Iberian collection contains a remarkably high genetic and phenotypic diversity, including adaptive variation in life-history traits (Picó *et al*., 2008; Méndez-Vigo *et al*., 2011; Marcer *et al*., 2018; Tabas-Madrid *et al*., 2018; Castilla *et al*., 2020) and the highest genomic diversity from the species’ native Eurasian range (The 1001 Genomes Consortium, 2016).

Here, we quantified the extent of chemical variation in root exudates across Iberian *A. thaliana* accessions by combining ultra-high performance liquid chromatography with quadrupole time-of-flight mass spectrometry (Subrahmaniam *et al*., 2023). We estimated broad-sense heritability values of root exudates, thereby assessing their degree of genetic determination. Furthermore, by conducting genome-wide association analyses, we also provided insight into the genetic basis of regional-scale variation in root exudates. Finally, we examined the eco-evolutionary forces putatively driving natural variation in root exudates by correlating chemical variation among accessions with their geographic, environmental, life history, and genetic patterns of variation across the Iberian Peninsula.

## Materials and methods

### Source accessions

The accessions included in this study are part of the Iberian *A. thaliana* collection, encompassing hundreds of populations from the southwestern Mediterranean Basin (Spain, Portugal, and Morocco; Picó *et al*., 2008; Brennan *et al*., 2014; Castilla *et al*., 2020). We sampled populations in annual field campaigns in spring from the early 2000s to late 2010s. In each field campaign, seeds from several individuals per population (around 6-7) were collected and multiplied using the single seed descent method in greenhouse conditions (Centro Nacional de Biotecnología, Madrid, ES) to increase seed number and quality while minimizing environmental and maternal effects. During these multiplication experiments, and when possible, we selected one maternal line per population based on average values per population for adaptively important life-history traits (e.g. flowering time). These maternal lines became accessions used for characterizing phenotypic traits and obtaining whole-genome sequences in subsequent studies (Tabas-Madrid *et al*., 2018). We kept multiplied seeds in cellophane bags under dry conditions, at room temperature, and in darkness for long-term storage. To avoid aging effects, seeds used in this study came from an additional multiplication experiment conducted in the late 2010s.

Previous genetic structure analysis on the Iberian *A. thaliana* collection revealed the existence of four genetic clusters: NW-C1, NE-C2, Relict-C3, and SW-C4 (cluster names as in Castilla *et al.,* 2020) (Supplementary Dataset 1). The three non-relict clusters are geographically structured as indicated by cardinal directions in their names, while the relict cluster is scattered mostly across the southern half of the Iberian Peninsula (Marcer *et al*., 2016). Evidence indicates that the relict cluster, which has several unique phenotypic and molecular features, probably endured dramatic environmental changes over millennia, representing part of the species’ early history (Durvasula *et al*., 2017; Toledo *et al*., 2020). The Iberian *A. thaliana* collection is also well characterized ecologically (Marcer *et al*., 2016; Vidigal *et al*., 2016; Tabas-Madrid *et al*., 2018; Castilla *et al*., 2020) (Supplementary Dataset 1), which is important for this study. In short, geographic coordinates and altitude of all accessions were recorded during field campaigns with a GPS (Garmin International, Inc., Olathe, US; positional error = 4 m). Climatic data associated to accession coordinates were extracted from WorldClim v.2 (Fick & Hijmans, 2017) and the Digital Climatic Atlas from the Iberian Peninsula (http://opengis.uab.es/wms/iberia/en_index.htm). Vegetation data came from the CORINE Land Cover 2000 (https://land.copernicus.eu/pan-european/corine-land-cover) as the percentage of vegetation types within a 500 m radius around the GPS coordinates of each accession. Finally, topsoil pH came from The Soil Geographical Database from Eurasia v.4 (https://esdac.jrc.ec.europa.eu/tags/soil-geographical-database-eurasia).

### Plant growth and root exudate sample collection

We initially selected 131 natural *A. thaliana* accessions from four genetic clusters, located in areas with less than 50% urbanized land (Marcer *et al*., 2016). Using a majority rule for genetic cluster memberships, the sample set included 91 accessions from NW-C1, 17 from NE-C2, 14 from Relict-C3, and 9 from SW-C4. This distribution of accessions reflects the natural abundance of *A. thaliana* in areas with low human influence for each genetic cluster. Clusters with a broad distribution, such as NW-C1, span diverse environments, whereas those with narrower distributions, such as SW-C4, are more specific to environmental conditions from smaller regions.

In July-September 2022, seeds from each accession were grown in five replicates at the Department of Molecular Biology and Genetics (Aarhus University, DK) using a protocol for handling large samples (Subrahmaniam *et al*., 2023). Briefly, all plants were grown *in vitro* at 21°C with a 16/8-hour light/dark cycle. Each Petri plate (total 153 plates) containing Murashige and Skoog medium with agar housed 4-5 accessions and one control. Plants were grown on autoclaved polypropylene mesh for facilitating easy removal. All plates were rotated periodically to minimize position effects. Col-0 was also used as a phytometer to gauge micro-environmental variation. All accessions were sampled six weeks after germination, during a specified period in the day (between 08:00 and 11:00 a.m.) to minimize diurnal variations. For exudate collection, each plant was removed using the meshes, roots were cleaned with a brush, and then transferred to 400 µL of ultrapure water for five minutes to capture erroneous exudates. Individual plants were then moved to 400 µL MilliQ water in 20 mL cylindrical glass vials, with roots submerged and the above-ground part floating with the help of the mesh. Vials were sealed to ensure sterility and trays containing vials were covered with aluminum foil at the base to simulate dark conditions. Trays were then placed in a growth chamber at 21°C for two hours. Afterwards, plants were removed from the vials and samples were immediately frozen in liquid nitrogen and stored at -18°C until further processing. Negative controls were prepared by treating empty vials following the same procedure.

### Root exudate chemical analysis

Overall, we retained 378 plant samples, comprising 3-4 replicates for 105 out of the 131 accessions, due to loss of plants during growth and/or the hydroponic root exudate sampling. Based on majoritarian genetic cluster membership, the final set of 105 accessions encompassed 75 accessions from NW-C1 (mean cluster membership proportion ± SE = 0.70 ± 0.01; range = 0.37–0.90), 10 from cluster NE-C2 (mean ± SE = 0.55 ± 0.03; range = 0.40– 0.70), 13 cluster Relict-C3 (mean ± SE = 0.77 ± 0.02; range = 0.64–0-88) and 7 from cluster SW-C4 (mean ± SE = 0.59 ± 0.03; range = 0.50–0.68). Some accessions exhibited mixed levels of membership consistent with some degree of admixture among genetic clusters (Fig. 1a), which provides a snapshot of the demographic and evolutionary complexity of *A. thaliana* across the region. It must be noted that the placement of accessions among clusters was not a biasing factor in this study design because we instead used genetic cluster proportional memberships in our analyses as a covariate (see below).

**Fig. 1.**
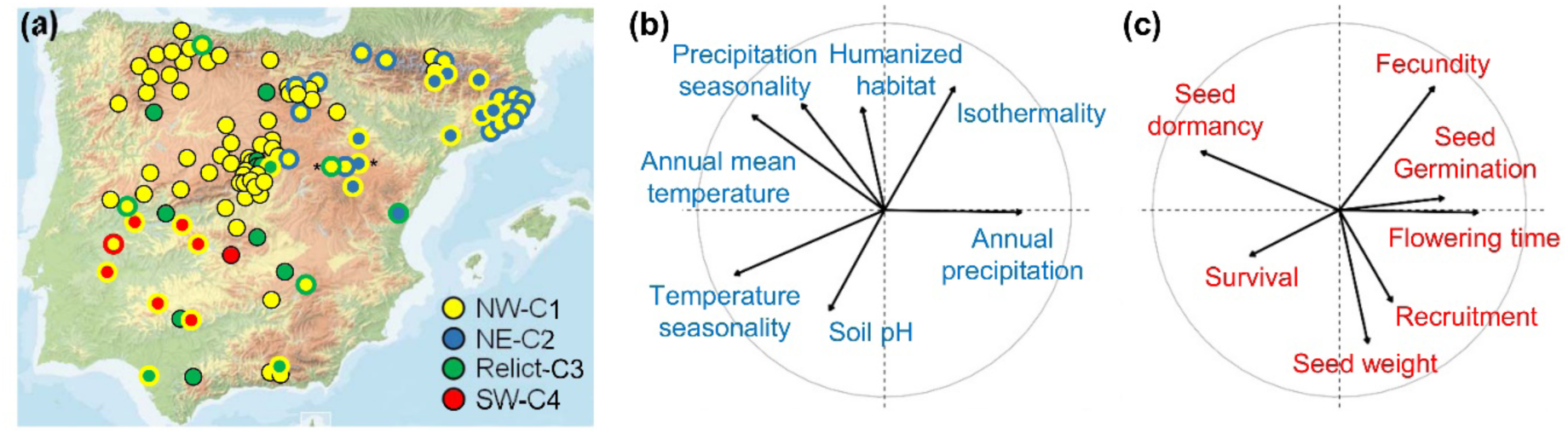
Geographic, genetic, environmental and life-history diversity of Iberian *A. thaliana* accessions used to estimate root exudate composition (see data in Supplementary Dataset 1). (a) Geographic distribution of 105 accessions across the Iberian Peninsula. Accessions are color-coded based on the assigned genetic cluster as follows. Dots with a unique color indicate accessions with proportional memberships higher than 0.5 in a single cluster and lower than 0.25 in the other three clusters. Dots with two colors indicate accessions with a predominant membership in a single cluster (inner color) and higher than 0.25 in another cluster (outer color). Only two accessions exhibited proportional memberships higher than 0.25 in three clusters (indicated by asterisks). In these two cases, we indicate the proportional memberships of the two main clusters following the same rationale as before. (b-c) Principal component analysis (PCA) plots of environmental variables (b) and life-history traits (c) characterizing all 105 accessions. Plots indicate the first two PCA axes and arrow length is proportional to the contribution of each variable/trait.

In September-December 2022, samples were subjected to untargeted metabolic profiling in negative and positive ionization modes using ultra-high performance liquid chromatography coupled with quadrupole time-of-flight mass spectrometry (UHPLC-QTOF-MS) at the Department of Chemistry of Aarhus University (Aarhus, DK). The UHPLC was an Ultimate 3000 (Thermo Scientific Dionex, Sunnyvale, US) and the QTOF-MS was a Bruker Compact (Bruker, Billerica, US). Instrument stability was assessed using a standard solution of six authentic standards (cinnamic acid, naringenin, L-tryptophan, quercetin, salicylic acid, and p-coumaric acid) injected throughout the run (Broadhurst *et al*., 2018). Additionally, an internal standard of ketopinic acid (10µg/mL in MilliQ water) was prepared for spiking all samples. Quality control samples were injected every 30 samples and biological control blanks every 20 samples. The m/z error averaged <3 ppm, retention time error was <1%, and peak area error was <10% in negative mode. In positive mode, only L-tryptophan was adequately ionized, with an m/z error <2 ppm, retention time error <1%, and peak area error <10%, indicating no significant instrumental drift throughout both runs.

However, quercetin, known to be photosensitive, exhibited greater variability in signal stability over time. Despite these fluctuations, the signals remained within the predefined warning and action thresholds, so that the variability was not due to analytical instability.

### Raw data processing

We processed the raw LC-MS data using MZmine 3 (Schmid *et al*., 2023) and SIRIUS 5 (Dührkop *et al*., 2021). MZmine 3 performs preprocessing, peak detection, alignment, quantification, and statistical analysis for untargeted metabolic profiling (Ludwig *et al*., 2018). Features were analyzed with SIRIUS 5, which uses isotopic pattern and fragmentation trees for structure elucidation and chemical annotation. Specifically, SIRIUS 5 uses CSI:FingerID and CANOPUS to predict molecular fingerprints and compound classes (Ludwig *et al*., 2020; Dührkop *et al*., 2021). CANOPUS integrates ClassyFire and Natural Product Classifier (NPC) to assign compound classes with a flexible three-tiered system (pathway, superclass, and class), allowing compounds to fit into multiple categories (Kim *et al*., 2021) while providing a Tanimoto similarity for structural matching (Ludwig *et al*., 2020). In evaluating the putative chemical compound prediction, a combination of SIRIUS chemical classes and Tanimoto scores was endorsed (Zulfiqar *et al*., 2023).

In total, we detected 1,142 features in negative ionization mode and 126 in positive mode across 105 Iberian *A. thaliana* accessions. Of these, 992 and 108 were identified as putative chemical compounds down to the NPC pathway level in negative and positive ionization modes, respectively, while up to 150 (∼13%) and 18 (∼14%) unidentified features were excluded in negative and positive modes, respectively. No features overlapped between negative and positive modes due to the high noise threshold imposed and the lower number of features obtained in positive mode.

### Data filtering for analysis

We converted absolute peak area data of features from MZmine 3 into relative data by dividing absolute peak area of a feature by the total peak area within the respective sample to estimate the contribution of each feature to the entire exudate composition of each sample. We excluded features present in only one replicate of any accession, eliminating 345 (∼30%) and 24 (∼19%) features in the negative and positive modes, respectively. Additionally, to avoid bias, features that were present in either less than 10% or more than 90% of the accessions— 470 (∼41%) in negative and 56 (∼44%) features in positive mode—were also omitted due to their rarity or prevalence, respectively. This filtering process resulted in 327 features in negative mode and 46 in positive mode for further analysis. Of these final 373 features, 358 (∼95%) were successfully annotated at the compound class level using CANOPUS, with 125 (∼33%) having a Tanimoto score >50%, displaying structural similarity to known compounds (Ludwig *et al*., 2012) (Supplementary Dataset 2). For clarity in subsequent sections, we refer to these 373 features as putative compounds as their identity has not been confirmed using authentic standards.

### Broad-sense heritability values

For each of the 373 compounds, we estimated the broad-sense heritability (*H^2^*) as *V_B_ / (V_B_+ V_E_)*, where *V_B_* is the among-accession variance and *V_E_* is the environmental variance (among individual replicates within each accession) for the relative compound abundance (Supplementary Dataset 2). To estimate *V_B_* and *V_E_*, we employed a linear mixed model framework where compound variation (response variable) was modeled as a function of accession identity (random effect). Restricted maximum likelihood (REML) estimation was used to obtain variance component estimates using the *lmer* function from the *lme4* R package (Bates *et al*., 2015). By fitting the model, we obtained estimates of the variance among accessions (*V_B_*) and the total variance (*V_B_+ V_E_*). To quantify uncertainty in *H^2^* estimates, we used a bootstrap approach in which resampling was conducted at the accession level. For each bootstrap dataset, a linear mixed-effects model was fitted and *H^2^* recalculated. We then used the bootstrap distribution of *H^2^* estimates to obtain 95% confidence intervals (using quantiles of the bootstrap distribution). We added a small constant value (1×10^-10^) to the relative peak area values for each feature individual measurements prior to analysis. This adjustment ensured that no values became exactly zero, as it could lead to computational challenges.

### Distance matrices and analyses

We estimated the Bray-Curtis chemical distance among the 105 accessions using the relative peak area of the 373 compounds with the *vegan* R package (https://CRAN.R-project.org/package=vegan). Chemical distance among accessions was estimated for all compounds and for each of the seven NPC pathway levels. NPC levels included 59 alkaloids, 108 amino acids and peptides, 88 carbohydrates, 46 fatty acids, 43 shikimates and phenylpropanoids, 9 terpenoids, and 5 polyketides. When analyzing NPC pathways, we excluded 15 compounds because they could not be assigned to any NPC pathway. We additionally computed the chemical distance among accessions based on the first three Principal Coordinate Analysis (PCoA) axes of their variation using the *stats* R package (R Core Team, 2021).

We estimated the Euclidian geographic distance among accessions using GPS coordinates of each accession (Supplementary Dataset 1) with PASSaGE v.2 (Rosenberg & Anderson, 2011). In the case of genetic distance among accessions, we used whole-genome sequences previously obtained from 174 Iberian *A. thaliana* accessions including 2.8 million of single nucleotide polymorphisms (SNPs) (Tabas-Madrid *et al*., 2018). Genetic distance was calculated based on the proportions of allele differences between pairs of accessions using all non-singleton SNPs with TASSEL v.5.2 (Bradbury *et al*., 2007). Additionally, for each genetic cluster, we computed a genetic cluster distance among accessions based on proportional memberships (estimated in Tabas-Madrid *et al*., 2018; Supplementary Dataset 1), resulting in four additional genetic distance matrices. This allowed us to test whether genetic cluster membership affected root exudate composition for all compounds and for compounds classified in each of the seven NPC pathway levels.

We assessed life-history distance among accessions using quantitative data for life-history traits obtained from previous studies (Manzano-Piedras *et al*., 2014; Vidigal *et al*., 2016; Marcer *et al*., 2018) that had pairwise correlation coefficients lower than 0.7. These traits included seed weight, seed dormancy (as quantified by days of seed dry storage required to reach 50% of germination; DSDS50), seed germination in growth chamber conditions at 22 °C, and life-history traits estimated in a common garden experiment, such as recruitment, survival, flowering time and fecundity (see experimental details in Manzano-Piedras *et al.,* 2014; Supplementary Dataset 1). Combining all phenotypic traits—that directly or indirectly affect fitness in *A. thaliana*—allowed us to relate the phenotypic adaptive attributes of each accession to its environmental conditions of origin. Thus, we first reduced the phenotypic space by conducting a Principal Component Analysis (PCA) on life-history traits and then computed the Euclidian life-history distance based on the first three PCA axes (accounting for 65.3% of the variation) with PASSaGE (Fig.1b).

Finally, we estimated the Euclidian environmental distance among accessions using the environmental datasets available with a similar procedure as for life-history distance matrix. In short, we first selected environmental variables with pairwise correlation coefficients lower than 0.7 and with a clear biological interpretation. Thus, we eventually considered annual mean temperature, isothermality, temperature seasonality, annual precipitation, precipitation seasonality, the proportion of humanized habitat around each accession, and topsoil pH (Supplementary Dataset 1). We then estimated the Euclidian environmental distance based on the first three PCA axes (accounting for 71.9% of the environmental variation) with PASSaGE (Fig.1c).

We explored the drivers of variation in chemical compounds among 105 *A. thaliana* populations by correlating pairwise chemical distance matrices with geographic, environmental, life history, and genetic distance matrices. We used standardized data (by subtracting the mean and scaling by the variance) in all analyses. Analyses were conducted with the whole dataset (373 compounds), the set of compounds with *H^2^*values significantly different from zero, and for each set of compounds falling into each NPC pathway levels. We conducted Mantel tests with a Bonferroni correction for multiple testing with *vegan*.

### Genome wide association analysis

We conducted genome-wide association (GWA) analyses to identify genomic regions underlying variation in chemical composition of root exudates in the Iberian *A. thaliana* collection. To this end, we conducted GWA analyses using the average relative peak abundances of chemical compounds for each accession as traits with a set of 1.5 million non-singleton SNPs previously developed for the Iberian *A. thaliana* collection that were functionally annotated (Tabas-Madrid *et al*., 2018; Arteaga *et al*., 2021). The SNP dataset was filtered to retain only those SNPs possessing a minor allele frequency (MAF) of at least five accessions in the 105 accessions (MAF ≥5%). Heterozygous calls in these SNPs were rescored to the major frequency allele and outliers from frequency distributions of traits were eliminated (0.6% of outliers in the final dataset), as they both reduce GWA statistical power (Atwell *et al*., 2010).

To improve the biological interpretation of the results, particularly when known genes were identified, we streamlined the initial list of 373 putative compounds to only those that received a probability score greater than 0.7 in the NPC pathway, superclass, and class categories. This refinement resulted in 109 putative chemical compounds being included in the GWA analyses from the families of NPC pathway classes (alkaloids: 6; amino acids: 36; carbohydrates: 23; fatty acids: 19; shikimates and phenylpropanoids: 22; and terpenoids: 2; polyketides: 1). GWA analysis was also conducted using the subset of compounds with *H^2^* values significantly different from zero.

For the GWA analyses, we employed a mixed linear model (MLM) implemented in TASSEL. To adjust for population structure, we included a genetic kinship matrix, and estimated from the proportion of shared alleles (Atwell *et al*., 2010), as a covariate. SNPs were genotyped as binary variables (0 or 1) and their associations with the root exudate data were assessed using a linear regression model. We applied a stringent significance threshold of –log_10_(P) = 7.5 to detect the most significant SNP associations. We used a previous gene ontology (GO) annotation enrichment on the Iberian collection (Tabas-Madrid *et al*., 2018) to identify genes detected by the GWA analyses based on the positions of significant SNPs, considering the two nearest flanking genes in the case of SNPs located in intergenic regions.

## Results

### Chemical variation of A. thaliana root exudates in the Iberian collection

We pinpointed 373 putative chemical compounds as root exudates from 105 Iberian *A. thaliana* accessions. In the negative ionization mode, we identified 327 compounds (range of molecular masses = 121.02–927.37 Da). In the positive mode, we identified 46 compounds (range of molecular masses = 130.05–530.98 Da). We determined the putative chemical identity and classification of 358 of these compounds with SIRIUS 5, with 124 (∼33%) having a Tanimoto similarity >50%. The NPC categorizations revealed primary metabolites, namely amino acids and peptides, carbohydrates, and fatty acids. Additionally, we detected secondary metabolite classes, including alkaloids, terpenoids, shikimates and phenylpropanoids, and polyketides. Overall, primary and secondary metabolites made up approximately 55% and 45% of the identified compounds, respectively (Fig. 2).

**Fig. 2.**
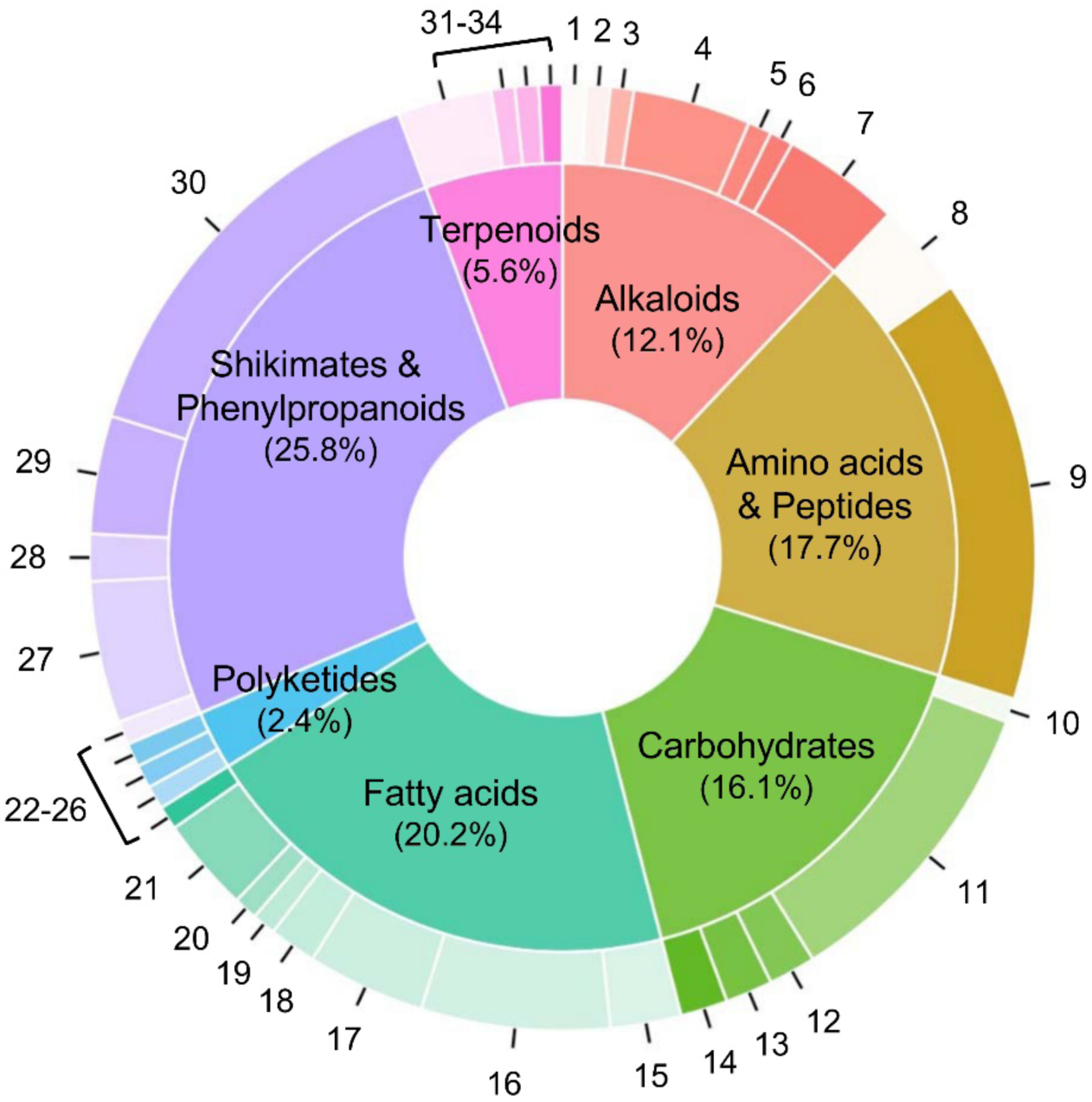
Donut chart displaying the distribution of putatively identified compounds in root exudates, categorized by the Natural Product Classification (NPC) pathway by superclass, in 105 Iberian *A. thaliana* accessions. The size of each inner segment corresponds to the number and percentage of compounds belonging to a specific pathway, whereas the size of each outer section represents the relative abundance of compounds within each specific superclass. Compounds (following the nomenclature provided by SIRIUS 5): 1: Amino acid glycosides; 2: Anthranilic acid alkaloids; 3: Nicotinic acid alkaloids; 4: Pseudoalkaloids; 5: Serine alkaloids; 6: Small peptides; 7: Tryptophan alkaloids; 8: Amino acid glycosides; 9: Small peptides; 10: Aminosugars/Aminoglycosides; 11: Nucleosides; 12: Polyols; 13: Saccharides; 14: Small peptides; 15: Fatty acids/Conjugates; 16: Fatty acyl glycosides; 17: Fatty acyls; 18: Fatty amides; 19: Fatty esters; 20: Monoterpenoids; 21: Octadecanoids; 22: Small peptides; 23: Macrolides; 24: Phenylpropanoids; 25: Polyethers; 26: Coumarins; 27: Flavonoids; 28: Lignans; 29: Phenolic acids; 30: Phenylpropanoids; 31: Apocarotenoids; 32: Lysine alkaloids; 33: Monoterpenoids; 34: Sesquiterpenoids.

We estimated broad-sense heritability estimates (*H^2^*) using relative abundance data for the 373 compounds across the 105 *A. thaliana* accessions. We identified 25 compounds with *H^2^* values significantly different from zero, ranging between 0.26 (95% CI = 0.0421-0.305) and 0.55 (95% CI = 0.310-0.664). Up to 24 of these compounds (range of molecular masses = 145.05–577.15 Da) were chemically categorized. These compounds were primary metabolites, such as amino acids and peptides (9) and carbohydrates (8), and secondary metabolites, such as shikimates and phenylpropanoids (3), alkaloids (2) and terpenoids (2).

The ordination analysis for the set of 373 compounds showed that the first two PCoA axes accounted for 28% of the variance in exudate chemical composition, where some separation between clusters was observed, particularly between Relict-C3 and SW-C4 (Fig. 3a). This distinction held for the second ordination analysis based on the 25 compounds with *H^2^* values significantly different from zero (Fig. 3b). In this case, the first two PCoA axes accounted for 56% of the variance in exudate composition (Fig. 3b).

**Fig. 3.**
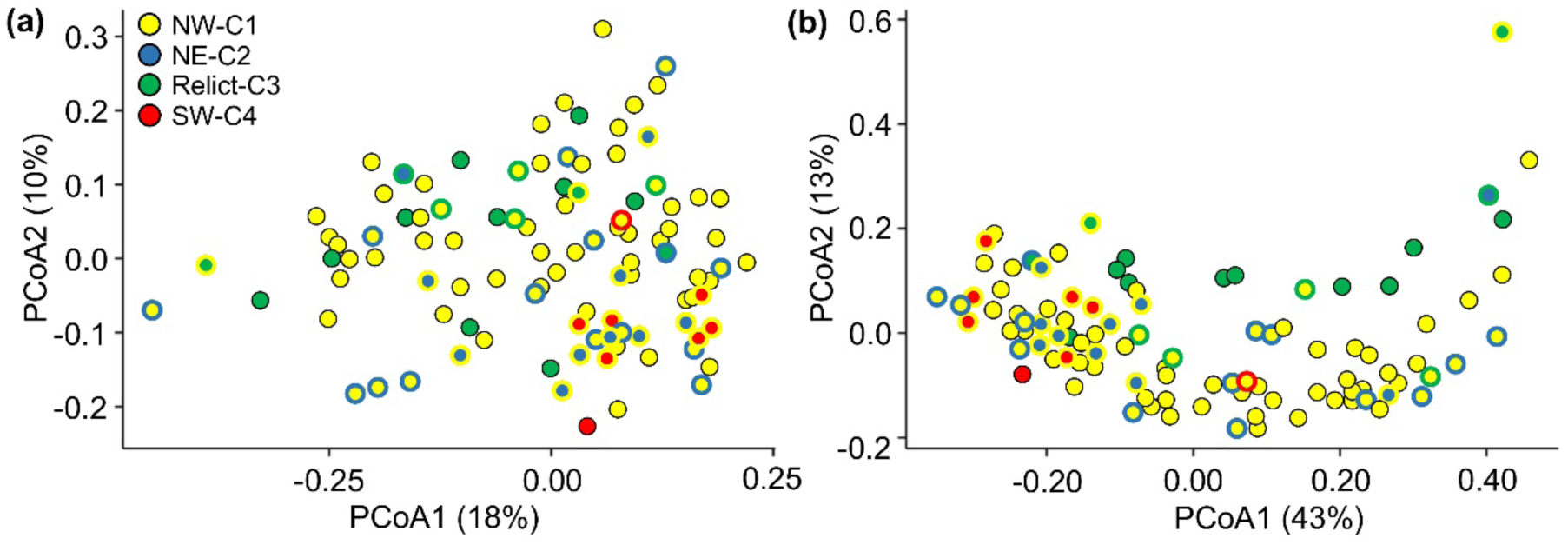
Principal Coordinate Analysis (PCoA) grouping 105 Iberian *A. thaliana* accessions based on root exudates. (a) Distribution of accessions based on 373 chemical compounds. (b) Distribution of accessions based on the 25 chemical compounds with broad-sense heritability values significantly different from zero. Accessions are color-coded based as in Fig. 1. The percentage of variation explained by the first two PCoA axes is given.

### Exploring eco-evolutionary associations with root exudate chemical variation

We found no significant correlations between chemical distances based on the 373 putative compounds and the 25-compound subset with *H^2^* values significantly different from zero, and any genetic (whole-genome distance), geographic, environmental, or life history distances (*P*>0.624 in all cases). We found the same result when compounds were categorized by their putative NPC pathways (*P*> 0.100 in all cases), except for a significant positive correlation between genetic distance and chemical distance for terpenoids (*r*=0.225, *P*=0.004).

The same set of analyses, but based on genetic distances using cluster proportional memberships, indicated that genetic clusters NE-C2 and SW-C4 did not show any significant relationships between genetic distance and chemical distance for either all compounds, or the 25-compound subset, or NPC category combination (*P*>0.072 in all cases). In contrast, genetic cluster Relict-C3 did show a positively significantly correlation between genetic distance and chemical distance for the 25-compound subset (r=0.100, *P*=0.032), meaning that accessions more similar genetically were similar chemically for this set of compounds. Interestingly, genetic clusters NW-C1 and Relict-C3 also showed positively significantly correlations between genetic distance and chemical distance for terpenoids (*r*=0.189 and 0.216, *P*=0.004 in both cases, for NW-C1 and Relict-C3, respectively).

We further evaluated terpenoid chemical differences between genetic clusters NW-C1 and Relict-C3 to detect those terpenoids further associated to their cluster proportional membership. To this end, we conducted one-way ANOVA testing the fixed effect of genetic cluster on abundance of each of the nine terpenoids detected using 1000 runs with down-sampled replicates (*n*=13) to avoid bias due to sample size differences between genetic clusters (*n*=13 and 75 for Relict-C3 and NW-C1, respectively). Up to five terpenoids exhibited significant differences between these two genetic clusters (*P*<0.035 in all cases; Fig. 4). In particular, compounds distyloside A, 8-epi-dihydro-penstemide, cyclolaudenol, and dinor-oxo-phytodienoic acid were present in NW-C1 and practically absent in Relict-C3, whereas atractyloside A was more abundant in Relict-C3 than in NW-C1 (Fig. 4).

**Fig. 4.**
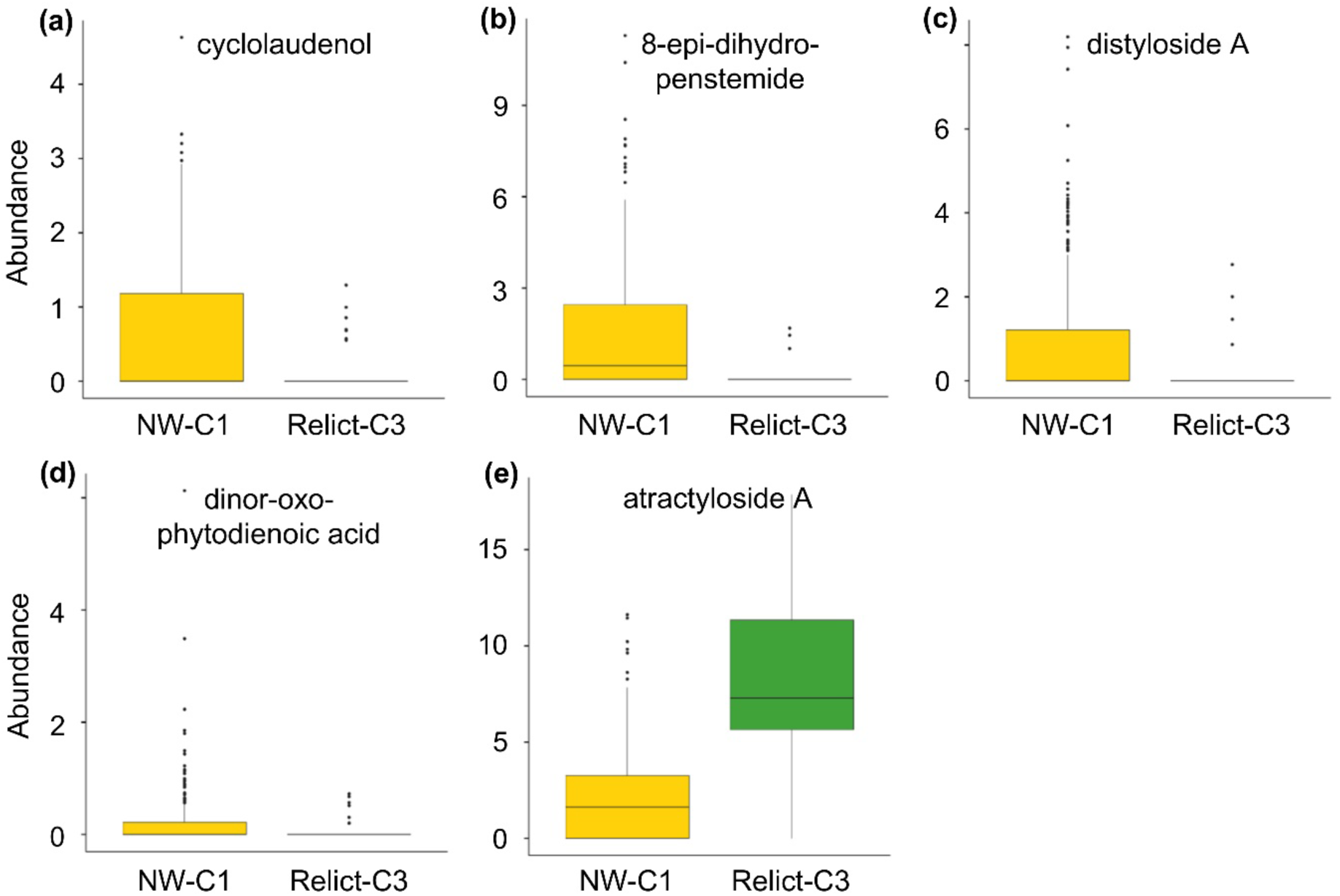
Variation in the abundance of five terpenoids found to be significant between genetic clusters NW-C1 and Relict-C3 in 105 Iberian *A. thaliana* accessions. Boxes show the lower and upper quartiles, whiskers are drawn down to the 10th percentile and up to the 90th, the line is the median of observations, and dots indicate data points falling out of percentiles. To obtain real values, numbers of axes need to be multiplied by 10^-3^ in all panels, except in panel d that need to be multiplied by 10^-2^.

### Genomic regions underlying chemical variation in root exudates

The GWA analyses identified 93 SNPs located in or nearby 70 genes that were significantly associated with 15 of the 109 compounds characterized in root exudates and eventually used in this analysis. Additionally, from the subset of 25 compounds with *H^2^* values significantly different from zero, we detected 37 SNPs linked to 10 genes. Further refinement of candidate genes based on biological pathway knowledge and gene ontology, led to the selection of 26 genes with known functional roles aligning with 12 compounds in our dataset, including 22 genes associated with nine compounds from the larger 109-compound dataset, and four genes linked to three compounds of the 25-compound dataset (Table 1). These genes were classified into four broad functional categories (i.e. metabolism, signaling and regulatory networks, nutrient transport, and stress response), although some genes fell in more than one category (Table 1).

**Table 1.** Genomic associations with root exudate chemical variation in Iberian *A. thaliana* accessions. The chemical family classification by SIRIUS 5, the specific chemical compound associated to each gene, the chromosome harboring significant SNPs, the position of the SNP in the chromosome (repeated positions indicate that SNPs were intergenic), the closest locus to the identified SNP, the protein encoded by the closest locus, and the functional classification describing the protein biological role are given. Functional classes include metabolism, signaling and regulatory networks, nutrient transport, and stress response. Superindexes 1 and 2 in the family column indicate the use of the 109-and 25-compound datasets, respectively, in GWA analyses. Asterisks next to compound codes indicate that Tanimoto scores for these compounds were >50%. Data sorted by chromosome number and position in the chromosome. Unknown proteins indicated by dashes.

| Family | Compound | Chr. | Position | Locus | Protein | Classification |
| --- | --- | --- | --- | --- | --- | --- |
| Fatty acids<sup>1</sup> | N.V1092 | 1 | 1497156 | AT1G05170 | B3GALT2 | Metabolism |
| Fatty acids<sup>1</sup> | N.V1092 | 1 | 1497156 | AT1G05180 | AXR1 | Signaling |
| Carbohydrates<sup>1</sup> | N.V919 | 1 | 10227720 | AT1G29265 | MIR399A | Nutrient |
| Fatty acids<sup>1</sup> | N.V195* | 1 | 11473826 | AT1G31950 | TPS29 | Metab/Stress |
| Carbohydrates<sup>1</sup> | N.V919 | 1 | 12264707 | AT1G33800 | GXM3 | Metabolism |
| Fatty acids<sup>1</sup> | N.V1092 | 1 | 17345313 | AT1G47317 | – | Stress |
| Alkaloids<sup>2</sup> | N.V831 | 1 | 21976138 | AT1G59740 | NPF4.3 | Nutrient |
| Alkaloids<sup>2</sup> | N.V831 | 1 | 21976138 | AT1G59740 | ARF1 | Signaling |
| Amino acids<sup>1</sup> | N.V313* | 1 | 22280140 | AT1G60470 | GOLS4 | Stress |
| Carbohydrates<sup>1</sup> | N.V554 | 1 | 22319649 | AT1G60600 | ABC4 | Nutrient |
| Amino acid<sup>2</sup> | N.V672 | 2 | 12605976 | AT2G29360 | SDR | Metabolism |
| Amino acids<sup>1</sup> | N.V313 | 2 | 18065356 | AT2G43500 | NLP8 | Nutrient |
| Amino acids<sup>1</sup> | N.V665* | 3 | 8721212 | AT3G24140 | FAMA | Signaling |
| Amino acids<sup>1</sup> | N.V665* | 3 | 8764452 | AT3G24220 | NCED6 | Metabolism |
| Amino acids<sup>1</sup> | N.V665* | 3 | 8769918 | AT3G24225 | CLE19 | Signaling |
| Amino acids<sup>1</sup> | N.V665* | 3 | 8778298 | AT3G24240 | RCH2 | Stress |
| Amino acids<sup>1</sup> | N.V665* | 3 | 8802982 | AT3G24290 | AMT1-5 | Nutrient |
| Amino acids<sup>1</sup> | N.V665* | 3 | 8808763 | AT3G24300 | AMT1-3 | Nutrient |
| Fatty acids<sup>1</sup> | N.V1092 | 4 | 2395964 | AT4G04720 | CPK21 | Stress |
| Shikimates<sup>1</sup> | N.V1001* | 4 | 8252678 | AT4G14340 | CK1 | Signaling |
| Shikimates<sup>1</sup> | N.V371 | 5 | 6349177 | AT5G19010 | MPK16 | Signaling |
| Shikimates<sup>1</sup> | N.V371 | 5 | 7697150 | AT5G23000 | RAX1 | Signaling |
| Shikimates<sup>1</sup> | N.V371 | 5 | 7714057 | AT5G23010 | MAM1 | Metab/Stress |
| Shikimates<sup>1</sup> | N.V371 | 5 | 7719834 | AT5G23020 | MAM3 | Metab/Stress |
| Carbohydrates<sup>2</sup> | N.V688 | 5 | 8389956 | AT5G24540 | BGLU31 | Metab/Stress |
| Amino acids<sup>1</sup> | N.V736 | 5 | 18367078 | AT5G45340 | CYP707A3 | Metabolism |

The metabolism-related genes included those involved in the synthesis of key cell wall polysaccharides, in particular, *B3GALT2* (putative beta-1,3-galactosyltransferase 2) and *GXM3* (glucuronoxylan 4-O-methyltransferase 3). Genes related to abscisic acid (ABA) synthesis included *CYP707A3* (abscisic acid 8’-hydroxylase 3) and *NCED6* (9-cis-epoxycarotenoid dioxygenase linked to putative t-butoxycarbonylleucylproline). Other genes involved in secondary metabolism were those related to terpenoid (*TPS29*; terpenoid cyclases linked to putative [10E,12E]-9,14-dioxooctadeca-10,12-dienoic acid), glucosinolate biosynthesis (*MAM1*, *MAM3*; methylthioalkylmalate synthase 1 and 3, respectively), alkaloid modification (*SDR*; tropinone reductase homolog), and phenolic compound modulation (*BGLU31*; beta-glucosidase 31).

The genes identified within the signaling and regulatory network category included transcription factors *FAMA*, *AUXIN RESPONSE FACTOR1*, *REGULATOR OF AXILLARY MERISTEMS1*, *AUXIN RESISTANT1*, *CLE19* (a CLAVATA3/CLE-related protein 19), *CK1* (casein kinase 1), and *MPK16* (mitogen-activated protein kinase 16). These genes are integral to various biological processes, such as cell fate determination, axillary meristem regulation, auxin signaling, cell-to-cell communication, cell cycle control, and mitogen-activated protein kinase signaling pathways. In this category, *FAMA* and *CLE19* were linked to putative glucobassicin, while *CK1* was linked to putative isorhamnetin 3-glucoside-7-rhamnoside.

Nutrient transport-related genes included *NPF4.3* (NRT1/PTR), a critical transporter for plant hormones; *MIR399A*, a microRNA essential for phosphate homeostasis; *NLP8* (NIN-like protein 8), involved in nitrogen metabolism signaling; *ABERRANT CHLOROPLAST DEVELOPMENT 4*, key in electron transport as a polyprenyltransferase; and *AMT1-3* and *AMT1-5*, which are important ammonium transporters. In this category, *NLP8* was linked with t-butoxycarbonylleucylproline, whereas *AMT1-3* and *AMT1-5* were linked to glucobrassicin.

In the stress response category, we detected genes, such as *GOLS4,* a galactinol synthase 4, *CPK21*, a calcium-dependent protein kinase 21, and *RGFR1*, a receptor-like protein kinase 2. These genes play significant roles in osmoprotection, innate immunity and calcium signaling. Notably, *BGLU31*, *MAM1*, *MAM3* and *TPS29* can also be grouped under this category for their secondary metabolite induced defense functions. Here, *GOLS4* was linked with t-butoxycarbonylleucylproline, while *RGFR1* was linked to glucobassicin.

## Discussion

Root exudates play several ecological roles that define a broad spectrum of plant responses to changes in the abiotic and biotic environment with which they interact. Notwithstanding their ecological and evolutionary value, the analysis of the amount and diversity of root exudates has been traditionally challenging for plant biologists for long-lasting technical reasons. Because of such limitations, our understanding of the genetic and environmental factors accounting for patterns of variation in root exudates at large geographical scales, which depicts major evolutionary trajectories in heterogeneous environments, has far remained unexplored. In this study, we bridged that gap of knowledge by providing the first regional-scale study of root exudate variation in the annual *A. thaliana* across the Iberian Peninsula, which enabled us to identify possible adaptive strategies related to biotic interactions that seem to be strongly conditioned by the genetic structure of accessions.

### Interpreting chemical complexity of A. thaliana root exudates

Our study identified large natural variation in chemical diversity of root exudates of *A. thaliana* across the Iberian Peninsula, including a wide range of primary (∼54%) and secondary metabolites (∼46%). Several primary compounds detected here were previously reported in *A. thaliana* root exudates, which may be especially critical to establish biotic interactions. For instance, malic acid is known to be involved in nutrient uptake and root-microbe interactions (Rudrappa *et al*., 2008; Ma *et al*., 2022), galactose and glucose affect rhizobacterial communities (Little *et al*., 2019; Lopes *et al*., 2022), whereas inositol plays a role in signal transduction and stress responses (Shears *et al*., 2012; Jia *et al*., 2019). Among the various amino acids and peptides detected, it is worth noting that these primary compounds in root exudates fluctuate with environmental and developmental stages (Chaparro *et al*., 2013; Strehmel *et al*., 2014). This is probably because they act as endogenous signals—influencing growth, development, primary metabolism, stress responses, and pathogen communication (Tavormina *et al*., 2015; Miller *et al*., 2017)—as well as long-distance signaling agents (Stegmann *et al*., 2017; Takahashi & Shinozaki, 2019).

As far as secondary metabolites are concerned, they may reflect the plant’s adaptive interactions with its environment, such as plant defenses and response in stress situations, as previously reported in *A. thaliana*. For instance, sinapate derivatives function as UV protectants and antimicrobial agents (Engels *et al*., 2012; Magdziak *et al*., 2020), glucopyranosides have allelopathic effects structuring the microbial community (Kimura *et al*., 2015; Upadhyay *et al*., 2022), and glucobrassicin and derivatives are well-known for plant defense against herbivores and pathogens (Madloo *et al*., 2019; Soengas *et al*., 2023). In addition, sinigrin is involved in water transport under salt stress, increasing hydraulic conductance and water permeability (Martínez-Ballesta *et al*., 2013). It must be emphasized, however, that our untargeted approach led to putative annotations with only one-third of the identified compounds having a Tanimoto confidence score above 0.5, which represents an important level of uncertainty in compound identification. While the methodology conducted in this study provided first insights into root exudate composition in *A. thaliana* at a regional scale, confirming and quantifying these metabolites require targeted analyses using authentic standards (Subrahmaniam *et al*., 2023).

### Genetic background of chemical variation in root exudate composition

This study underscored the notorious natural variation in root exudate chemistry of Iberian *A. thaliana*. It is worth emphasizing that, due to technical limitations, previous studies explored genotypic variation in *A. thaliana* root exudate composition in rather limited sets of accessions (9 worldwide accessions; Houshyani *et al*., 2012) and other genetic materials (19 MAGIC parental lines; Mönchgesang *et al*., 2016). Here, we vastly increased sample size up to 105 natural accessions across the Iberian Peninsula that enabled us to identify up to 373 putative chemical compounds. Such a step forward to quantify root exudate diversity also represented an opportunity to assess the genetic background of root exudates in *A. thaliana* in two complementary ways.

First, we were able to assess broad-sense heritability (*H^2^*) values of every root exudate, which is evolutionary important because they provide an estimate of the degree of genetic determination, that is, the proportion of phenotypic variance explained by genotypic variance. As experiments conducted in a common environment minimize the environmental component of phenotypic variation among accessions, adaptive processes are assumed to be an important source of variation in traits with *H^2^* values significantly different from zero. The results showed that only 25 of 373 compounds (6.7%) exhibited *H^2^* values significantly different from zero, primary and secondary metabolites constituting 67% and 33% of these 25 compounds, respectively. The greater proportion of primary metabolites with *H^2^* values significantly different from zero aligns with the accepted idea that natural selection is stronger on primary metabolites, which are fundamental in cell biological functions, and more relaxed on secondary metabolites, which are further related to environmental interactions (Kooke & Keurentjes, 2012). Nevertheless, the low proportion of compounds with *H^2^*values significantly different from zero also indicates that variation in root exudates is mostly influenced by environmental cues. Given the important and multiple direct or indirect functions of root exudates in all sort of plant responses to abiotic and biotic environmental interactions, which are per se extremely variable across space and probably over timescales, the massive proportion of both primary and secondary compounds with *H^2^* values indistinguishable from zero clearly reflects their environmentally driven nature.

Second, the availability of whole-genome sequences for all Iberian *A. thaliana* accessions allowed us to explore the genomic architecture underlying regional root exudate variation, that is, to detect genomic loci statistically associated to regional-scale variation in root exudates by GWA analysis. We detected significant associations between polymorphic loci and root exudate variation in 12 compounds, identifying up to 26 genes associated to a wide spectrum of plant functions, including all major categories, such as metabolism, signaling, nutrient transport, and stress response/defense (Table 1). Although traits with high heritability normally led to the increase of statistical power for detecting causal variants in GWA analysis (Khanzadeh *et al*., 2021), we only detected four of 26 genes associated to root exudates with *H^2^*values significantly different from zero, pinpointing the complexity of the genetic attributes of root exudates. It is worth noting, nevertheless, that our GWA analysis must be considered as exploratory because we did not have any candidate gene to confirm, as occurs when conducting GWA analysis on *A. thaliana* traits with a strong adaptive value and a well-known genetic basis, such as flowering time (Tabas-Madrid *et al*., 2018).

### Eco-evolutionary drivers of natural variation in A. thaliana root exudates

We assessed the role of geography, environment, life history, and genetics as major drivers of variation in chemical differences in root exudates among Iberian *A. thaliana*. Except genetics, all drivers showed non-significant associations with root exudate variation, indicating that macro-environmental drivers of phenotypic variation normally used to characterize *A. thaliana* accessions across broad geographical scales cannot account for variation in root exudate composition. Despite the important environmental component of phenotypic variation detected in root exudate composition, our results suggest that environmental drivers of variation might be operating at the micro-environmental scale, which means that micro-environmental parameters of the spots from where accessions were sampled (e.g. vegetation type, soil composition, temperature/humidity patterns) might better characterize such drivers of variation. Clearly, further research is needed to identify the environmental drivers of root exudate variation and the precise micro-spatial scale at which they might be operating.

Turning to the genetic drivers of root exudate variation in Iberian A. *thaliana*, we only detected a significant relationship between genetic distance, based on whole-genome sequences, and chemical distance for terpenoids. Perhaps because terpenoids, which are a modified class of terpenes, mediate important below-ground interactions between plants and other organisms, such as microbes, herbivores, and other plants (Huang & Osbourn, 2019), they emerge as the only NPC category of root exudates with a clear association with genetic similarity among *A. thaliana* accessions. The relevance of terpenoids detected in this study is in line with a recent study on root exudate compositions using 65 plant species, where overall metabolome composition lacked a phylogenetic signal, but only phenol content in root exudates showed evidence for evolutionary conservatism (Rathore *et al*., 2023). On top of that, it must be emphasized that one of the genes detected in our GWA analysis, i.e. *TPS29*, is key for terpenoid synthesis within the isoprenoid pathway and is highly expressed in roots (Yu et al., 2020). The compound associated to *TPS29* is a fatty acid, i.e. (10E,12E)-9,14-dioxooctadeca-10,12-dienoic acid (Tanimoto score = 0.74), and both terpenoids and fatty acids originate from acetyl-CoA (Sasaki & Nagano, 2004), directly linking *TPS29* to fatty acid metabolism.

Further support on the importance of terpenoids in natural root exudate variation of Iberian *A. thaliana*, and their relationship with genetic attributes, was found in the significant associations between chemical distances and genetic distances using genetic cluster membership proportions. Once again, only variation in terpenoids appeared to be significantly associated to genetic distances for genetic clusters NW-C1 and Relict-C3, suggesting that genetic structure also matters to account for natural variation in terpenoid exudates in *A. thaliana*. Interestingly, four terpenoids in root exudates appeared to be significantly more abundant in accessions with higher proportional memberships to NW-C1 cluster, whereas only one clearly dominated in accessions with higher proportional memberships to Relict-C3 cluster (Fig. 4). As Iberian clusters depict different evolutionary and demographic histories (Picó *et al*., 2008; Marcer *et al*. 2016; Tabas-Madrid *et al*. 2018), such differences in specific terpenoids probably reflect specific evolutionary trajectories associated to each cluster. In particular, most of these terpenoids are known to be associated to plant defense in various ways, such as acting as mitochondrial inhibitors against herbivores (atractyloside A; Woyda-Ploszczyca, 2023), displaying antimicrobial and anti-inflammatory effects (8-epi-dihydro-penstemide; Zajdel *et al*., 2012), deterring herbivores and pathogens through its potential toxicity (distyloside A; Kim *et al*., 2011), and participating in defense signaling (cyclolaudenol; Ezzat *et al*., 2016).

These findings suggest that NW-C1 and Relict-C3 Iberian clusters might have evolved specialized chemical defenses in response to its particular ecological challenges across their broad and heterogeneous Iberian environments. We cannot interpret the implications of these findings without falling into excessive speculation about the role of the terpenoids identified in clusters NW-C1 and Relict-C3, or lack thereof in clusters NE-C2 and SW-C4. However, our results are particularly interesting for Relict-C3. In particular, recent studies indicated that Iberian relict *A. thaliana* exhibits unique traits not found in any other accession from all over the species’ range, such as trichomes on pedicels and fruits (Arteaga *et al*., 2021, 2022), that might have also evolved as defensive barriers against a wide range of herbivores (Fürstenberg-Hägg *et al*., 2013). Hence, the convergence of specific physical (fruit trichomes) and chemical (atractyloside A) defenses in Iberian relict *A. thaliana* suggests that interactions with herbivores/pathogens might represent an important evolutionary driver, on top of the association of this cluster with a more pronounced Mediterranean climate and non-humanized environments (Marcer *et al*., 2016; The 1001 Genomes Consortium, 2016; Toledo *et al*., 2020). Finally, the evolutionary relevance of atractyloside A (Tanimoto score = 0.65) received further support from the fact that this compound was also the one with the highest *H^2^* value significantly different from zero (*H^2^* = 0.55), stressing the high genetic determination of this compound in Iberian *A. thaliana* and therefore its putative adaptive value. Overall, this study calls for further research on the sort of biotic interactions of relict *A. thaliana* in natural environments to understand the development of such an array of defensive strategies.

## Conclusions

Our study generated the greatest description of natural variation in root exudates known to date, identifying 373 putative compounds in 105 Iberian *A. thaliana* accessions fully characterized environmentally, genetically, and phenotypically. Interestingly, our results strongly suggest an important effect of genetic structure on root exudate composition, particularly for terpenoids, which are mostly involved in plant defenses against herbivores and pathogens. Hence, our study stresses the evolutionary importance of biotic interactions mediated by root exudates in *A. thaliana*, which aligns with other studies highlighting the role of specific *A. thaliana*’s defense metabolites, such as glucosinolates, to understand adaptive variation mediated by biotic interactions at large geographical scales (Katz *et al*., 2021). Our study represents a first step towards unravelling the enormous diversity and ecological roles of root exudates, in particular for terpenoids, which emerged as the candidate compounds to look at more carefully in the future. Nevertheless, further research needs to be conducted to improve our understanding of root exudates, including quantification of root exudates at different developmental stages (e.g. vegetative *vs* reproductive), environments (e.g. controlled *vs* natural), and stress situations (e.g. hydric, thermic, pathogenic).

## Supporting information

Accession and compound data are available as Supplementary Datasets.

## Acknowledgements

We thank Prof. Simona Radutoiu, Dr. Alen Trgovcevic, and Raffaella De Masi from the Plant Molecular Biology group at the Department of Molecular Biology and Genomics of Aarhus University for guidance on *A. thaliana* cultivation. We also thank Master’s students Antigoni Christofili and Jonas Dahl Dueholm from the Department of Chemistry for their assistance in chemical analysis and sample preparation. We are grateful to Dr. Carlos Alonso-Blanco (CNB-CSIC) for his advice on the GWA analysis.

## Authors’ contributions

HJS, MG and BKE conceived the idea. HJS, FXP, TB, MG, CLS and BKE generated materials and data. HJS, TB, BKE and FXP analyzed the data. HJS wrote the first version of the manuscript and all authors contributed to it.

## Availability of data and materials

Data deposited in the Dryad repository: https://doi.org/10.5061/dryad.vmcvdnd2j (Subrahmaniam *et al*., 2024). Accession and compound data are available as Supplementary Datasets. Seeds from all accessions are publicly available through the Nottingham Arabidopsis Stock Centre (NASC; http://arabidopsis.info). Raw whole-genome sequences are available from previous publications (The 1001 Genomes Consortium, 2016; Tabas-Madrid *et al*., 2018). Field sampling was conducted in locations where permission was not required. Based on the Royal Decree of the Spanish legislation (Real Decreto 124/2017, de 24 de febrero; https://www.boe.es/eli/es/rd/2017/02/24/124), the genetic resources included in this study fall within the definition of “taxonomic purposes”.

